# Acute activation of autophagy enables growth plate regeneration following radiation-induced injury

**DOI:** 10.64898/2026.08.24.746001

**Authors:** Yashar Mehrbani Azar, Amal Nazaraliyev, Mahtab Avijgan, Lars Sävendahl, Klas Blomgren, Phillip T Newton

## Abstract

**Purpose:** Radiation injury to growth plates commonly leads to skeletal late complications including short stature, limb length-discrepancy, and scoliosis/kyphosis in pediatric oncology patients. We aimed to understand the acute responses of direct growth plate irradiation that result in skeletal late complications.

**Materials and methods:** We first established an *in vivo* model of focal growth plate irradiation that recapitulates the clinical development of skeletal late complications and used it to explore the responses of growth plate chondrocytes within the first 72 hours of radiation exposure. To monitor acute effects of radiation exposure on human chondrocytes, rare human growth plate biopsies were exposed to ionizing radiation *ex vivo*. Using these approaches, we applied clonal genetic tracing and immunofluorescence to monitor changes at the cellular and molecular levels. Functional *in vivo* perturbations were conducted with clinically-relevant autophagy inhibitor, hydroxychloroquine.

**Results:** Growth plate irradiation disrupted the continuous production of chondrocytes required for bone elongation and was associated with DNA damage throughout the growth plate. Indicators of growth plate activity, SOX9 and the phosphorylated form of ribosomal protein S6, decreased during a 6- and 24-hour post-irradiation window but returned to normal levels 72 hours after irradiation. We identified a surge in autophagic flux throughout the growth plate during this window, based on temporal SQSTM1 and LAMP1 protein levels. The earliest stages of these response mechanisms are conserved between species and relevant to humans. Hydroxychloroquine treatment immediately after radiation injury in mice impaired growth plate regeneration, resulting in more severe late complications.

**Conclusion:** Our findings demonstrate that autophagy is an important acute response to irradiation in growth plate chondrocytes, revealing a novel potential therapeutic target for preventing radiation-induced skeletal late complications.

## Introduction

Skeletal growth during childhood and adolescence is vital for humans to reach their final height and expected body symmetry. Since radiotherapy can damage the cartilage organs - called (epiphyseal) growth plates - that enable bones to grow in length, understanding the underlying mechanisms of damage and repair can facilitate better treatment options and reduce long term complications.

Growth plates are present near the ends of elongating bones and contain cartilage cells (chondrocytes) at various differentiation stages layered histologically in distinct zones [1,2]. The resting zone (RZ) contains slowly dividing stem and progenitor cells that are highly quiescent [3]. Offspring from the RZ chondrocytes give rise to stacks of proliferating zone (PZ) chondrocytes (ref., Fig. 1A). Once they stop dividing, the chondrocytes then enlarge to form the hypertrophic zone where extracellular matrix mineralization occurs [4]. Subsequently, most hypertrophic chondrocytes undergo cell death; this process gradually lengthens the bone as the space vacated by the dying cells becomes part of the underlying primary spongiosa - newly forming bone [5]. This growth mechanism requires the continuous production of new growth plate chondrocytes to maintain bone elongation. Previous studies in rodents have determined that irradiation of the growth plate leads to an immediate decrease in cell cycle progression, which resumes several days later [6].

**Figure 1.**
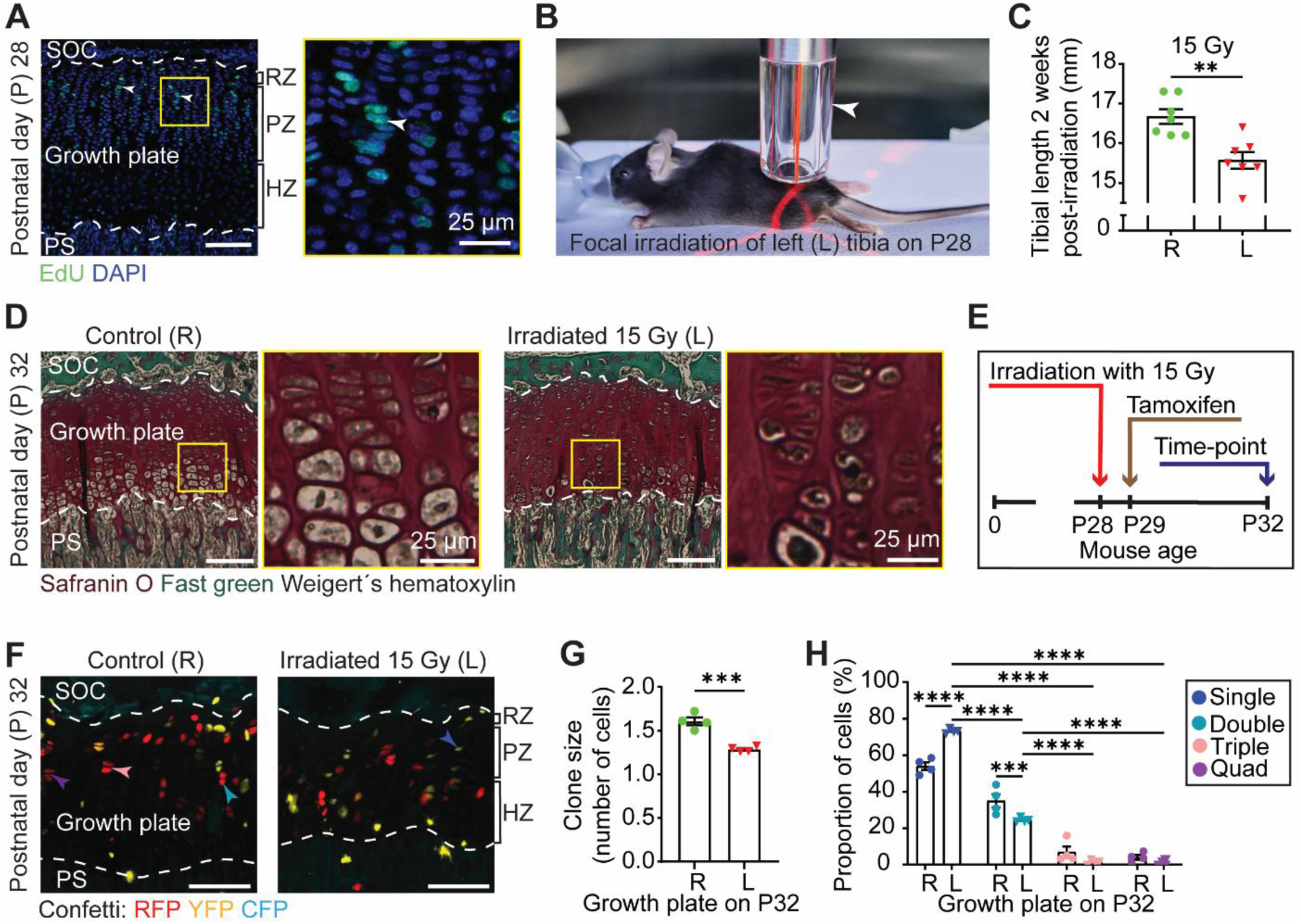
Ionizing irradiation rapidly impaired chondrocyte proliferation in the postnatal growth plate. (**A**) Proximal tibial growth plate from mouse at postnatal day 28, labelled with EdU 4 h before tissue collection to visualize proliferating chondrocytes (arrowheads). (**B**) Experimental setup of focal irradiation of the left (L) proximal tibial growth plate at postnatal day 28 under isoflurane anesthesia, where irradiation is applied in a field of 1 cm diameter using a perspex-tipped applicator (arrowhead). (**C**) Tibial length is reduced two weeks after irradiation with 15 Gy irradiation (L) compared to contralateral control (Right, R) (n=7 mice). (**D**) Histological images of proximal tibial growth plates on P32, four days after irradiation. Tissue sections were stained with Safranin O (red, cartilage), Fast Green (green, bone and connective tissue) and Weigert’s hematoxylin (black, nuclei). (**E**) Experimental set-up for clonal genetic tracing of irradiated chondrocytes using Col2CreERT:R26R-Confetti mice. (**F**) Clonal dynamics were analyzed in Col2CreERT:R26R-Confetti mice as per Fig. 1E and sections visualized by confocal microscopy. (**G**) Irradiation reduced average clone sizes (n=4 mice) and (**H**) the proportion of chondrocytes in clones of two (double), three (triple) or four (quad) cells. Scale bars = 100 μm, unless otherwise specified. The secondary ossification center (SOC), growth plate (GP), primary spongiosa (PS), and the resting (RZ), proliferating (PZ) and hypertrophic (HZ) zones are indicated for orientation, and the growth plate is demarcated by dashed lines.

At the interface between proliferative and hypertrophic zones, chondrocytes undergo enlargement coinciding with inhibited (macro)autophagy, hereafter referred to as autophagy [7]. Autophagy, deriving from the Greek, “self-eating” is a process occurring in virtually all cells to shuttle damaged or unwanted material, including macromolecules and organelles, to lysosomes for degradation [8]. Due to the protective effects autophagy can mediate, it has become a target in cancer treatment and post-irradiation repair [9].

Every year, approximately 250,000 children worldwide are diagnosed with cancer, and approximately a third of them receive radiotherapy [10]. While growth plates are not typical radiotherapy targets, it is often challenging for radiation oncologists to design irradiation dose plans that entirely avoid them. Direct irradiation damage to growth plates damages chondrocytes [11] causing skeletal late complications ranging from a loss of final height to severe skeletal malformations (such as scoliosis/kyphosis) that can cause painful, debilitating consequences to childhood cancer survivors [12–14]. With improved survival rates, increasing life expectancies and limited treatment options, childhood cancer survivors are affected by these conditions throughout their lives [15].

Here, we set out to model skeletal late complications *in vivo* and explore the acute effects of irradiation on growth plates that eventually lead to them.

## Materials and Methods

### Ethical approval and consent to participate

All experiments in this study were approved and performed in accordance with the guidelines from the Swedish National Board for Laboratory Animals and the European Union Directive (2010/63/EU) under an ethical permit (DNR 16673/2020) approved by the Stockholm’s Animal Experimentation Ethics Committee (Stockholms djurförsöksetiska nämnd). The collection of human growth plate cartilage was performed under an ethical permit (97/214) approved by the Stockholm Research Ethics Committee North.

### Study design

In all experiments, 3R principals were considered and the E-estimator (resource allocation equation) tool was used to determine sample size, with remaining mice within experimental litters typically included. Where mice needed to be selected for experiments from a pool, the ID numbers of the mice were randomly allocated into experimental groups by using the “List Randomizer” tool at random.org where a pre-determined position on the randomly generated list was used to assign animals to each group. Single-blinded analysis was performed when manual quantification was involved (Fig. 1G and H and Fig. 2D) by anonymizing the sample to the observer. An exception was made only for comparing groups with obvious changes where the observer could easily distinguish the groups based on their appearance. Both male and female mice were used in this study. Individual mice were considered as one experimental unit. A study design protocol was not registered prior to the onset of the project. The study was performed in accordance with the Animal Research: Reporting of *In Vivo* Experiments (ARRIVE) guidelines [16].

**Figure 2.**
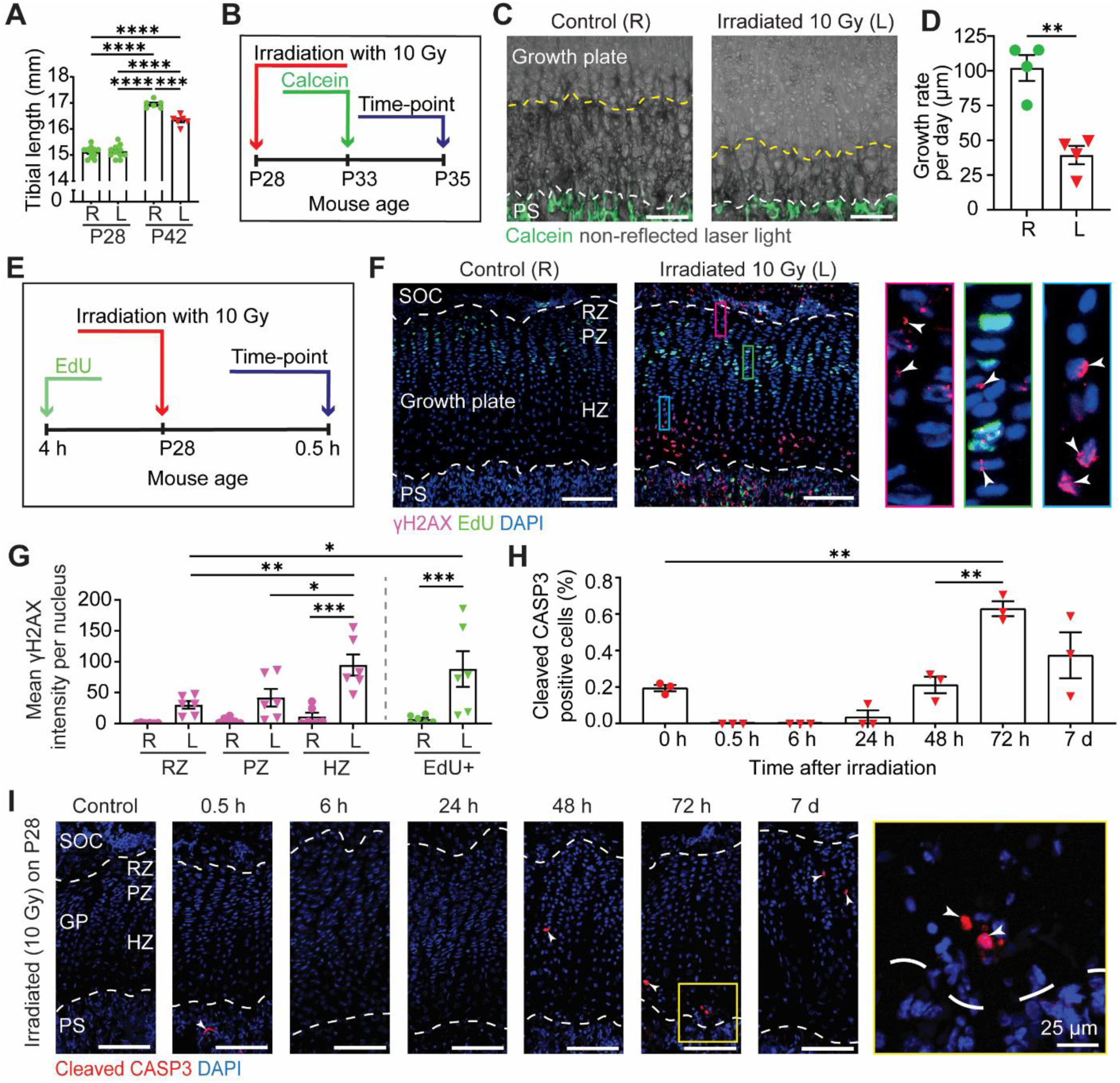
Focal tibial irradiation *in vivo* induced skeletal late complications after acute DNA damage within the postnatal growth plate. (**A**) Tibial length on P28 (n=11 mice; i.e. prior to irradiation) and on P42 (n=6 mice; fourteen days after irradiation of L tibia with 10 Gy). (**B**) Experimental set-up to measure growth rate after irradiation. (**C**) Images of mouse proximal tibial growth plate on P35, 48 h after calcein injection. (**D**) Quantification of daily growth rate (µm) with the mineralized front indicated with a yellow dashed line (n=4 mice). (**E**) Experimental set-up used to analyze DNA damage to growth plate chondrocytes. (**F**) DNA damage in dividing cells was visualized by combining EdU detection and immunofluorescence for γH2AX. (**G**) Quantification of γH2AX signal intensity per nucleus was performed (n=6 mice). (**H**) Percentage of cleaved CASP3-positive growth plate chondrocytes was quantified up to seven days after 10 Gy focal irradiation on P28. (**I**) Representative image of cleaved CASP3 immunostaining. Scale bars = 100 μm, unless otherwise specified. The secondary ossification center (SOC), growth plate (GP), primary spongiosa (PS), and the resting (RZ), proliferating (PZ) and hypertrophic (HZ) zones are indicated for orientation, and the growth plate is demarcated by dashed lines.

### *In vivo* irradiation procedure

All mice were irradiated (X-ray) using a CIX-3 cabinet (Xstrahl, Surrey, United Kingdom). During irradiation, mice were under isoflurane (Läkemedelsverket, QN01AB06) anesthesia (at 5 % induction and 2 % maintenance rate) with an air/oxygen ratio of 1:1. Anesthetized mice were placed in the prone position and irradiation was applied to the left knee of mice within a circular field of 1 cm diameter at a delivered rate of 1.260 ± 0.004 Gy/min (dosimetry uncertainty ∼ 2%; 300 KV and 10 mA) to target the proximal tibial growth plate, with the right side used as a contralateral control. This approach enables growth plate irradiation whilst avoiding damage to organs in the axial regions of the body, e.g., brain, liver, and digestive tract, that could indirectly affect bone growth. By postnatal day 28 (P28), the growth plate is fully formed, and fueled by stem cells [17]. An Xstrahl Perspex-tipped applicator was used to collimate the beam. External filtration giving a half-value layer (HVL) of 4.0738 mm Cu was applied by adding a Thoraeus filter (1.0 mm Sn, 0.25 mm Cu, 1.50 mm Al). Dosimetry was performed annually by Karolinska Institutet’s radiation physicist. Biologically effective doses (BEDs) were calculated as per the linear quadratic formula [18], using an alpha/beta ratio previously established for growing cartilage [19].

Depending on the experimental design, a single dose of 10 Gy or 15 Gy (exclusively in Fig. 1) was applied on postnatal day 28 (P28). Mice were given a single dose of tamoxifen (Sigma-Aldrich, T5648) on postnatal day 29 (P29) by oral gavage of up to 200 μg per g body weight, diluted (53.84 mM) in corn oil (Sigma-Aldrich, C8267) for experiments with the Col2CreERT:R26R-Confetti strain (Fig. 1F-H). Tibiae were carefully dissected from surrounding soft tissues and separated from the femur and foot. Tibial length was measured *ex vivo* using a digital caliper (Marketlab, 45072) as the distance between the most proximal and distal ends of the bone.

### *In vivo* hydroxychloroquine (HQ) treatment procedure

Hydroxychloroquine (Sigma-Aldrich, H0915) administration was performed in combination with unilateral growth plate irradiation. Mice received a single 10 Gy dose of X-ray irradiation to the left knee on postnatal day 28 (P28). Hydroxychloroquine was administered by oral gavage at a dose of 50 mg/kg (prepared fresh in sterile water) at two defined time points after irradiation. The first dose was delivered 6 h post-irradiation on day 28 (P28), and the second dose was administered 24 h after irradiation day 29 (P29). Control mice received equivalent volumes of vehicle.

### Mice strains

The Col2-creERT strain (developed by S. Mackem, US national institute of health [NIH]) and Rosa26R-Confetti (The Jackson Laboratory, 017492) were crossed for clonal studies, when the recombination [43,44] led to inexpression of Cre enzyme (Cre negative) then these mice were used in other experiments if possible (in line with the 3R principles). Mice from the commonly used C57BL/6J strain (The Jackson Laboratory, 000664) were utilized throughout the study in experiments related to Figures 2-6 in time points P28 control, 24 h and 72 h. Mice of both sexes were used in all experiments, with a total of 53 % male and 47 % female mice overall.

### Mice housing and husbandry

Mice were housed in Komparativ Medicin Biomedicum (KM-B) facility at Karolinska Institutet. Cages were individually ventilated, and mice had access to chow and water *ad libitum*. Litters were housed with their parents and siblings until weaning, at approximately three weeks of age, after which time they were separated and caged with their siblings of the same sex. Mice were group-housed with the parent mouse on a 12 h light-dark cycle at 22°C with 50% humidity. Animals were not handled by tail-lifting, tunnel lifting was used as standard.

### Genotyping

Genotyping was performed from biopsies (Mouse ear collected at weaning to identify positive recombination of *Cre* expressing Col2-creERT strain). For clonal genetic tracing, the R26R-Confetti strain was crossed with Col2-creERT mice to produce the Col2-creERT:Confetti strain.

In the Col2-creERT strain, developed by S. Mackem, US national institute of health (NIH), expression of CreERT protein is controlled by Col2a1 [1]. Cre-mediated DNA recombination is dependent on tamoxifen in this strain. Rosa26R-Confetti (The Jackson Laboratory, 017492, referred to as R26R-Confetti or Confetti) [44] is a reporter mouse strain that contains the brainbow 2.1 construct [45]. Upon Cre-mediated DNA recombination, one of four different fluorescent proteins (nuclear green, cytoplasmic red, cytoplasmic yellow and membrane-bound cyan) is expressed in a stochastic manner from each allele, allowing clonal identification [44].

DNA extraction per sample: Tissue samples were lysed in 75 µl lysis buffer (25 mL of 100 mM NaOH; 0.4 mL of 50 mM Na2EDTA (pH 8); 74.6 mL of NFW [nuclease free water]) over a heat-block (95°C) for thirty minutes. Then they were neutralized by adding 75 µl neutralizing buffer (pH 5) (40 mM Tris-HCl; in 74.6 mL of NFW) and vortexed.

PCR (Polymerase chain reaction) per sample: Primers *Cre* reverse (Merck, SY200208702-054 [5’-CCCACCGTCAGTACGTGAG]) 10 µM and *Cre* forward (Merck, SY200208702-053 [5’-CGCGGTCTGGCAGTAAAAA]) 10 µM were premixed. PCR Master mix was prepared (Gotaq G2 Green Master Mix [Promega, 7823] 12.5 µl; NFW 11 µl; Primer pre-mix 1 µl) and 1 µl of each sample was added to 24 µl of master mix. Tubes were placed in thermocycler (Applied Biosystems, 2720 thermal cycler) and chain reaction continued (1 cycle: 94°C 5 min; 33 cycles: 94°C 30 sec, 63°C 6 sec, 61.5°C 6 sec, 60°C 6 sec, 60.5°C 6 sec, 60°C 6 sec, 59°C 6 sec, 72°C 45 sec; 1 cycle: 72°C 7 min; Hold 4°C).

Electrophoresis and imaging: 1.7% agarose (Meridian Bioscience Agarose Molecular Grade, BIO41025) in 100 ml TAE (Tris-acetate-EDTA) was prepared containing 4 µl Gel Red (Biotium, BIT-41003). Gel was placed in the electrophoresis container (BioRad, Power Pac Basic) filled with TAE and 10 µl of each sample (of 25 µl PCR reaction) was Loaded to each chamber. Electrophoresis was run at up to 130 V for the desired period. Bands in gel were visualized by Fusion imaging box (Fusion FX Vilber Lourmat, Ramcon) and Fusion software (FusionCapt Advance Fx7 17.03) using auto-exposure.

### Tissue processing for mouse samples

Mice were euthanized by cervical dislocation under isoflurane anesthesia and both hind-legs were collected. Bone lengths were measured using digital calipers (Thermo Fisher Scientific, 11777105). For the preparation of frozen sections, the dissected tissues were placed in glass vials (Millipore Sigma, V7130) containing freshly prepared pre-cooled (4°C) formaldehyde 4% (Histolab, HL96753.1000) in 0.1 M PBS (phosphate-buffered saline) (Thermo Fisher Scientific, Gibco™ PBS, 10010056) and were fixed for 6 h at 4°C on a roller depending on experiment. The formaldehyde solution was then replaced with 30% sucrose (Nordic Biosite, 105-S-3333) and rolled again overnight at 4°C. Then tissues were embedded in OCT compound (Sakura Tissue-Tek, 4583) in Cryo Mold (Sakura Tissue-Tek, 4557 [25 x 20 x 5 mm]) and after freezing over dry ice, stored at -20°C. Sections of between 30 μm and 150 μm in thickness were prepared with a cryostat (Leica, CM3050). Sections were mounted over SuperFrost Ultra Plus™ GOLD adhesion slides (Thermo Fisher Scientific, K5800AMNZ72) and were kept at -20°C.

For the preparation of paraffin-embedded sections (used only in Fig. 1D), dissected tissues were immediately fixed in freshly prepared, pre-cooled (4°C) 4% formaldehyde in 0.1 M phosphate-buffered saline (PBS) for 48 h with gentle rocking. Samples were briefly washed with PBS at 4°C, then decalcified with 10% ethylenediaminetetraacetic acid (EDTA) tetrasodium salt dihydrate (Scharlau, AC09671000) in Milli-Q water, adjusted to pH 8.05, at room temperature with gentle rocking. The EDTA solution was changed every two to three days. After decalcification, samples were stored in 70% ethanol (EtOH) until further processing. Tissues were processed for dehydration and embedded in paraffin wax (Milestone, LOGOS). Sections of 5 µm thickness were collected using a microtome (Thermo, HM355S) on SuperFrost Ultra Plus™ GOLD adhesion slides.

Growth plates were analyzed using longitudinal sections through the proximal tibia. To ensure consistency between samples, tissues were sectioned through the central region of the knee joint using the cruciate ligaments as an anatomical landmark. This approach minimized variation between samples and ensured that comparable regions of the growth plate were evaluated across all experimental groups.

### Preparation and irradiation of human growth plate samples

Human growth plate samples were collected from constitutionally tall-stature patients (Table 1) undergoing epiphyseal surgery (epiphysiodesis) to reduce bone growth at Karolinska University Hospital [46]. Immediately after collection, the biopsies were transferred into DMEM (Dulbecco’s Modified Eagle Medium)/F-12 medium (phenol red free [Gibco, 21041025]) on ice and transported directly to the laboratory. Biopsies in DMEM/F-12 were prepared into slices approximately 1 mm thick with a scalpel under dissection microscopy on ice. Each human growth plate slice was placed in an individual well of a 24-well plate (Sarstedt, 83.3922) in 2 ml of culture medium consisting of DMEM/F12 medium (Gibco, 11320033) supplemented with 50 μg/ml ascorbic acid, 0.2% BSA (Sigma-Aldrich, 05470-5 G), and 50 μg/ml gentamicin (Fisher Scientific, 11530506). The wells containing the control biopsies were shielded with lead (3 mm aluminum filter) and ionizing X-ray radiation was applied to the 24-well plate using a single dose of 2 Gy on a 40 FSD rotating platform (195 mV, 10 mA). External filtration giving a half-value layer (HVL) of 4.0738 mm Cu was applied by adding a Thoraeus filter (1.0 mm Sn, 0.25 mm Cu, 1.50 mm Al). The irradiation was delivered at a rate of 1.260 ± 0.004 Gy/min. Dosimetry was performed annually by Karolinska Institutet’s radiation physicist. After irradiation, samples were cultured in a humidified atmosphere (37°C, 5% CO2) for 6 h and then fixed in pre-cooled 4% formaldehyde/PBS for 6 h at 4°C. Samples were then placed into 30% sucrose (AG Scientific, S-2885) overnight and embedded in OCT medium in cryomolds. Samples were stored at -80°C prior to cryo-sectioning. Sections of 30 μm thickness were prepared with a cryostat (Leica, CM3050) and on SuperFrost Ultra Plus™ GOLD adhesion slides and were kept at -20°C.

**Table 1.** Metadata of the patient information for the tissues analyzed in. Fig. 5.

| Patient ID | Genetic sex | Tanner pubertal stage B<sup>1</sup> | Tanner pubertal stage G<sup>2</sup> | Patient age at surgery |
| --- | --- | --- | --- | --- |
| 1 | M | - | 3 | 13 y 11 m |
| 2 | F | 3 | - | 12 y 11 m |
| 3 | M | - | 3 | 13 y 10 m |

### Safranin O and Fast Green staining

Slides containing 5 µm paraffin-embedded sections were rehydrated for 5 min in each panels containing (xylene [Fisher Chemical, X51], xylene, EtOH 99%, EtOH 95%, EtOH 70%) and stained with Weigerts hematoxylin (Merck, H3136) dipped in Acid Alcohol solution (1% HCl (Merck, 1.43007.1000):70% ethanol) and washed in distilled water. Slides were then stained with 0.02% Fast Green solution (Merck, F7258) followed by a 1% acetic acid solution then, without rinsing, 1% Safranin O (Merck, S8884) solution. Slides were rinsed in 95% ethanol and dehydrated (EtOH 95%, EtOH 99%, EtOH 99%, xylene, xylene) 5 min in each, before mounting. Images (Fig. 1D) were captured by a bright field (10X optical) microscope (Axio Imager).

### Labelling with Calcein

Mice were injected intraperitoneally with Calcein (Merck, C0875) 10 μg per g body weight, 48 h before tissue collection. The samples were collected, placed into 4% formaldehyde for 6 h at 4°C on a roller, tissue processing was as described above. Slide preparation and mounting: few drops of PBS gently applied and covered the slides to remove OCT at room temperature, then PBS was drained, and slides were covered with anti-diffuse fluorescence mounting medium (Fluoroshield™) (Sigma-Aldrich, F6182-20ML) at room temperature, coverslips (VWR, 631-0147) were placed over slides. Images were captured by confocal microscope and the ImageJ software (NIH, Bethesda, U.S.A.) was used to measure the distance between mineralization front and the metaphyseal side of the growth plate (Fig. 2C).

### EdU labelling and detection

EdU (Life Technologies, E10187) was injected intraperitoneally 4 h before irradiation at concentration of 65 μg per g body weight. After sample preparation (30 µm tissue sections), EdU was detected using a click reaction for 30 min at room temperature in a mixture containing 0.1 M tris aminomethane (Tris, pH 7.5) (Sigma-Aldrich, 10812846001), 2 mM CuSO4 (Sigma-Aldrich, 451657), 0.1 M ascorbic acid (Sigma-Aldrich, A5960) and 2 μM Alexa Fluor azide 647 (Invitrogen, A10277). Slides were mounted with Fluoroshield™ and imaged (Fig. 1A and 2F) by confocal microscope. Quantification was conducted using ImageJ.

### Immunofluorescence protocols

Samples were cryosectioned (Leica BIOSYSTEMS [CM3050 S], Danaher, USA) at 30 µm thickness and mounted on Superfrost Plus Gold slides (Epredia, FT4981IGLPLUS-001), which were stored at -20°C. Frozen slides were thawed at room temperatures for 10 min, a humidified chamber was prepared, and the OCT was washed from slides by covered PBS for 15 min. Antigen retrieval was achieved as follows: no retrieval (anti-phospho-RPS6, anti-γH2AX and anti-SQSTM1), 0.1% trypsin in PBS(T) (0.05% Tween 20 [Sigma-Aldrich, P9416-100ML] or 0.05% Triton X100 [Sigma-Aldrich, T8787-100ML]) for 30–45 min at room temperature (anti-SOX9, anti-LAMP1 and anti-cleaved CASP3). Then slides were Washed three times for 5 min with PBS(T) 0.05% Triton X100. After blocking with 3-5% horse serum (Jackson ImmunoResearch, 008000121) in PBS containing 0.1-0.3% Triton X100 for 30–60 min, slides were incubated with primary antibody: anti-cleaved CASP3 (Cell Signaling Technology, 9579, 1:250); anti-LAMP1 (Santa Cruz Biotechnology,19992, 1:250); anti-SOX9 (Merck, AB5535, 1:500); anti-phospho-RPS6 (Cell Signaling Technology, 4858S, 1:100); anti-γH2AX (Cell Signaling Technology, 9718, 1:200); anti-SQSTM1/p62 (PROGEN, GP62-C, 1:500) overnight at 4°C (very gentle shaking), followed by three times wash for 5 min with PBS(T) (0.05% Tween 20) and subsequently incubated with the appropriate secondary antibody: Alexa Fluor 647 (Thermo Fisher Scientific, A-31573, 1:800); Alexa Fluor 555 (abcam, ab150154, 1:500); Alexa Fluor 555 (Thermo Fisher Scientific, A-31572, 1:400); Alexa Fluor 488 (Jackson ImmunoResearch Laboratories, 706-545-148, 1:500) for 60–90 min at room temperature over shaker under protection from light. Followed by counterstaining (20 min) with DAPI (4’,6-diamidino-2-phenylindole, dihydrochloride) (4.28 mmol) (Sigma-Aldrich, D9542, 1:500), and again washed three times with PBS after. Slides were then mounted with Fluoroshield™.

### Microscopy and image analysis

Confocal imaging (Carl Zeiss, LSM700, Oberkochen, Germany) equipped with ZEN software (Blue edition, Carl Zeiss) was used for clonal analysis and after immune staining as described previously (Zhou *et al.*, 2019). Briefly, Z-stack images were acquired in sequential scans using a 20X objective lens. The images were displayed as maximum intensity projection, unless otherwise stated. Contrast, brightness and gamma adjustment have been altered to improve visualization in pseudo-colored confocal scans; where needed, unirradiated control samples stained alongside experimental samples were used to guide visualization parameters. Neighboring tiles were automatically stitched together by the ZEN software. Histochemical images were captured by bright field microscope (Axio Imager M2; Carl Zeiss microscopy, Germany), equipped with the Stereo Investigator software (Micro Bright Field Inc.).

### Quantification of fluorescent signal

ImageJ software was used for the digital analysis of confocal images. The Freehand Selection tool was selected to outline the growth plate and outside the line was cleared. All the spaces surrounding the growth plate were filled in black color. For direct comparison of the signal between samples, images were stitched together in a single file (noted order) leaving a gap between each image and groups. Every pixel was selected, and the signal was converted to a graph using the plot profile tool. The Wand tool was used to quantify the area under the curve for each plot, and the data were exported to an Excel file, where each value representing fluorescent signal was normalized by the number of chondrocytes (marked by DAPI) in the same area of the corresponding growth plate.

### Retrieval of clinical trial data

Data obtained from clinicaltrials.gov on 3^rd^ February 2025 is presented in Table 2. Search terms “everolimus”, “chloroquine”, “3-methyladenine”, “sirolimus” and “hydroxychloroquine” were applied. Studies with eligibility of “Child”, defined as individuals between 0 and 17 years of age, were included. Trials were deemed active when status was listed as “Recruiting”, “Not yet recruiting” or “Unknown”. Some spacing and punctuation changes were made to the text for presentation purposes.

**Table 2.**
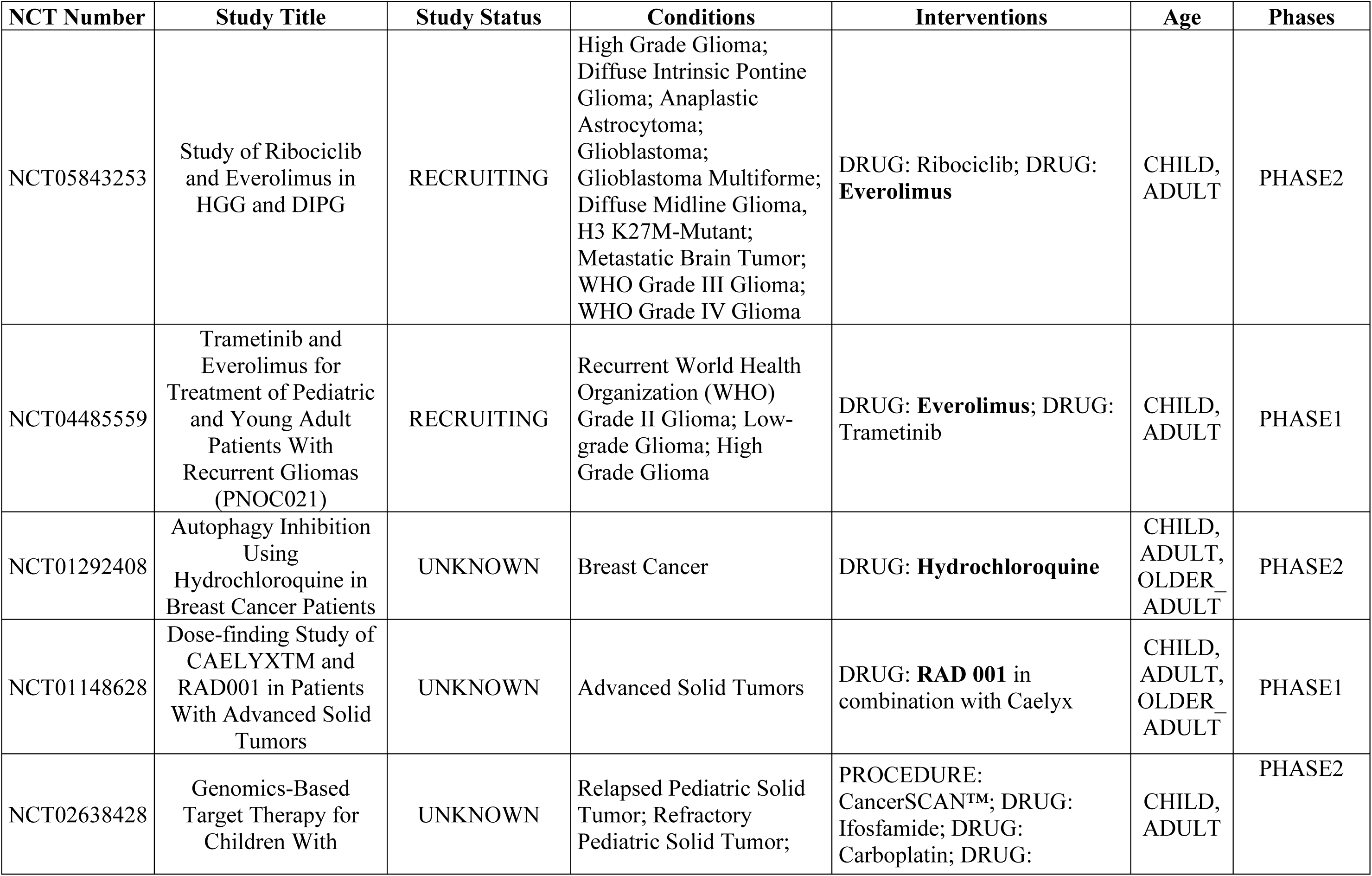
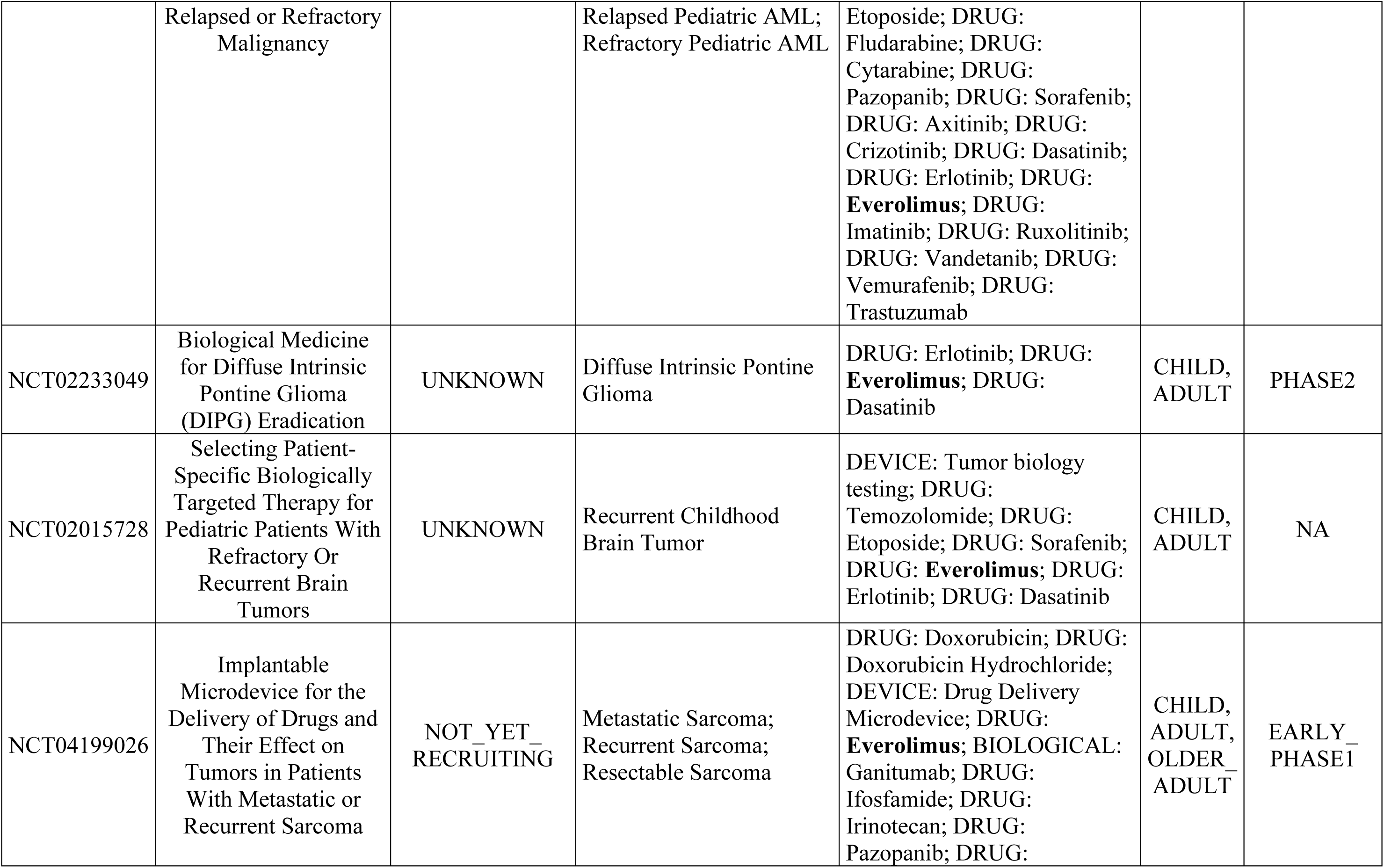
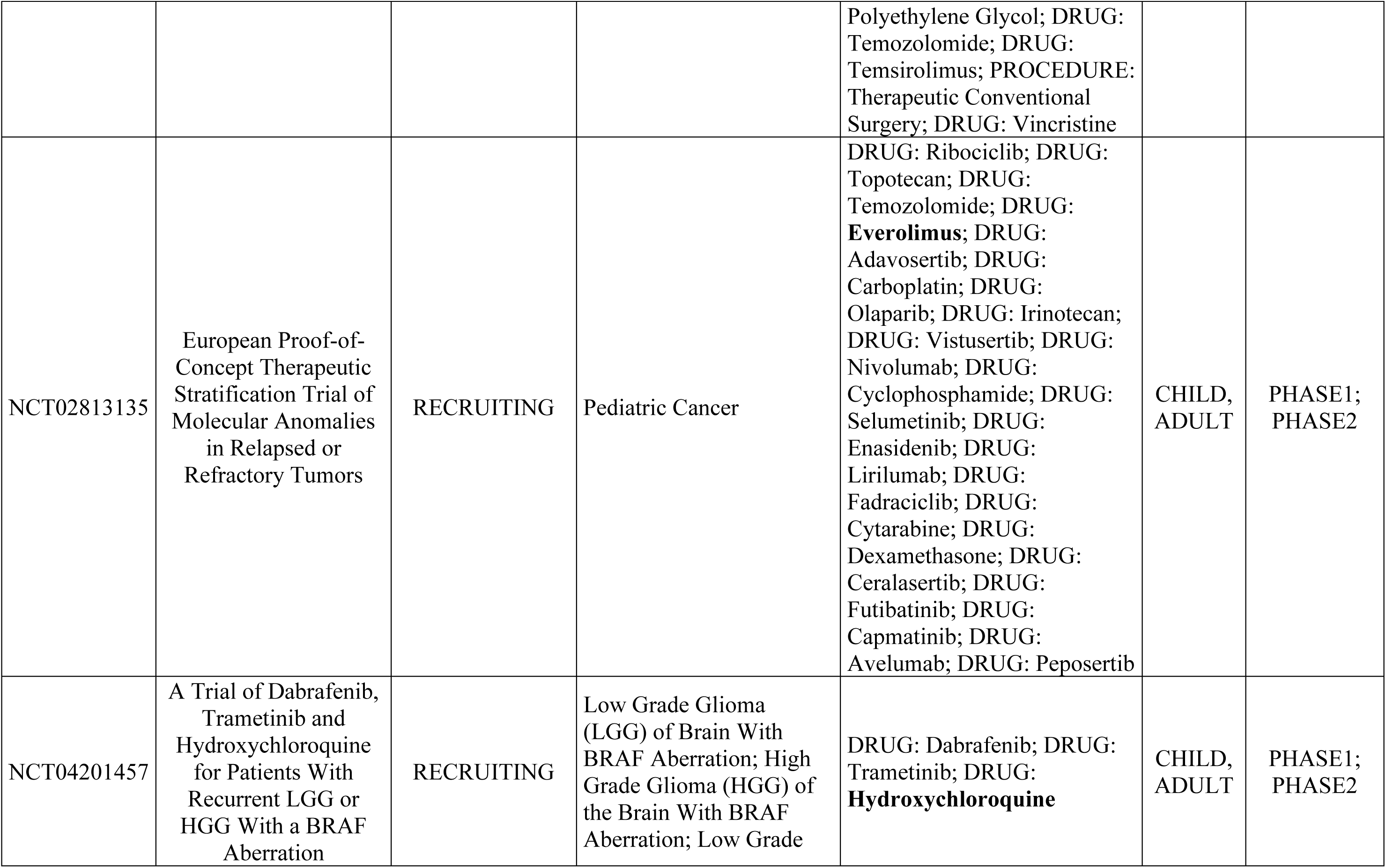
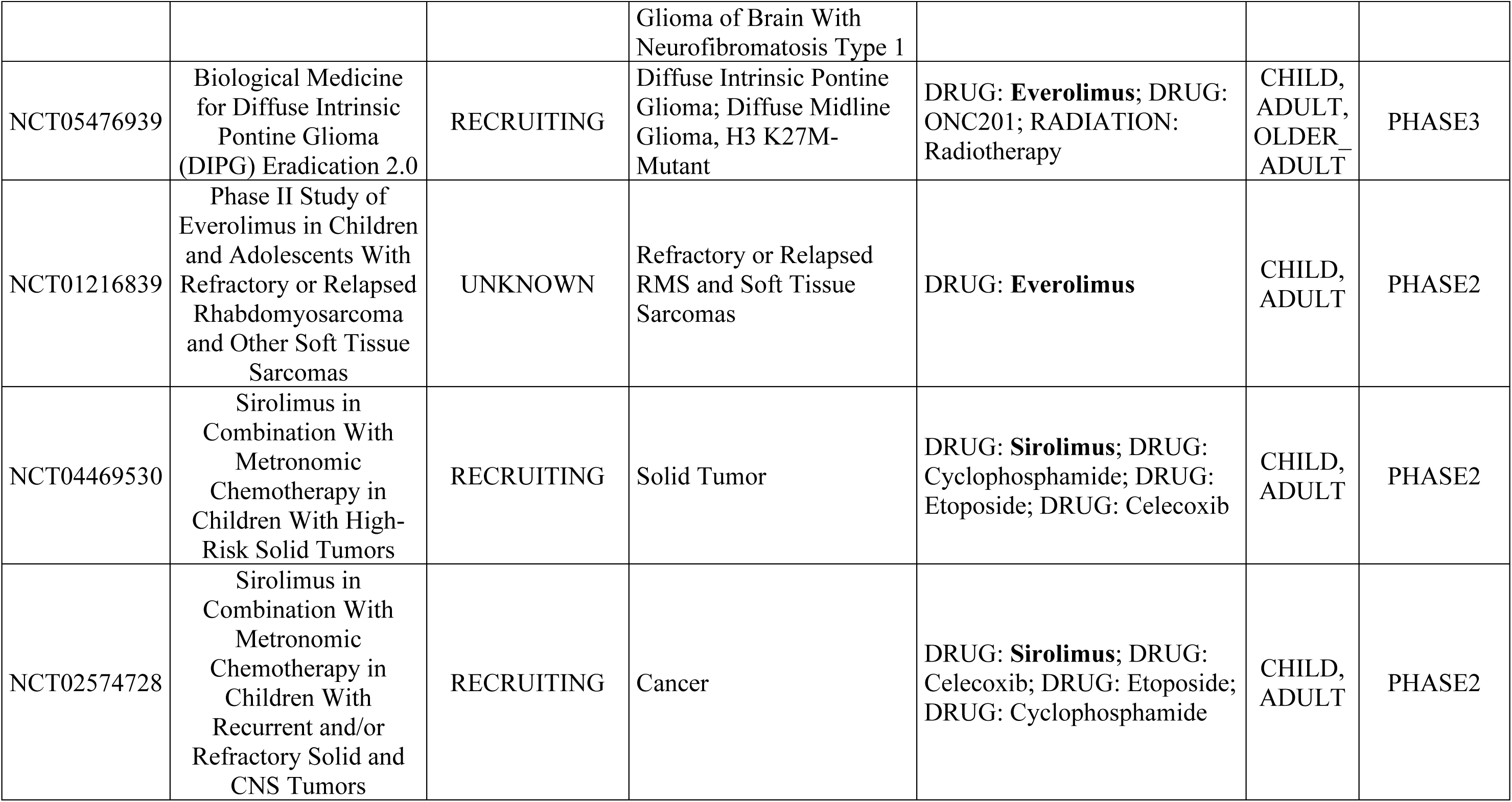
A summary of ongoing clinical trials in which autophagy modulators are being tested in pediatric oncology patients. The names of autophagy modulators have been highlighted in bold text.

### Software

Figures and graphics (in Fig. 5A) were produced using Adobe Illustrator V. 29.2.1 (Adobe, San Jose, California, United states). Specific software are mentioned in specific methods.

### Statistics

All quantifications were presented as individual data-points with the mean ± standard error of the mean (SEM), unless otherwise indicated. Outliers were identified as values that were outside of two standard deviations of the mean, which were excluded prior to analysis. The Shapiro-Wilk test for normality was performed before each analysis to test for normality. When data fulfilled normality criteria, unpaired Student’s t-test was used to compare two groups (6B, 6D, 6F, 6H, 7C, 7E-F), except when comparisons between irradiated and contralateral limbs originated from the same animal, when a paired t-test was used (Fig. 1C, 1G and 2D), and one-way ANOVA with Tukey’s multiple-comparison test was used to calculate P-values when more than two groups were compared (Fig. 2A, 3C, 3D, 3I, 3J, 4C, 4D and 4G-I), unless otherwise specified. If the analysis did not pass the normality test, then the non-parametric Mann-Whitney test was used to compare two groups and Kruskal-Wallis test with Dunn’s multiple comparison was applied to determine statistical significance (Fig. 2H, 3E, 3H and 4B). For Fig. 1H and Fig. 2G, two-way ANOVA with Bonferroni’s multiple comparisons test was performed. All statistical analyses were carried out using Microsoft Excel and GraphPad Prism version 10.3 (GraphPad Software, La Jolla, California, United States). P-values are presented as *p<0.05, **p<0.01, ***p<0.001, and ****p<0.0001 throughout. Statistical differences were considered significant when p≤0.05.

**Figure 3.**
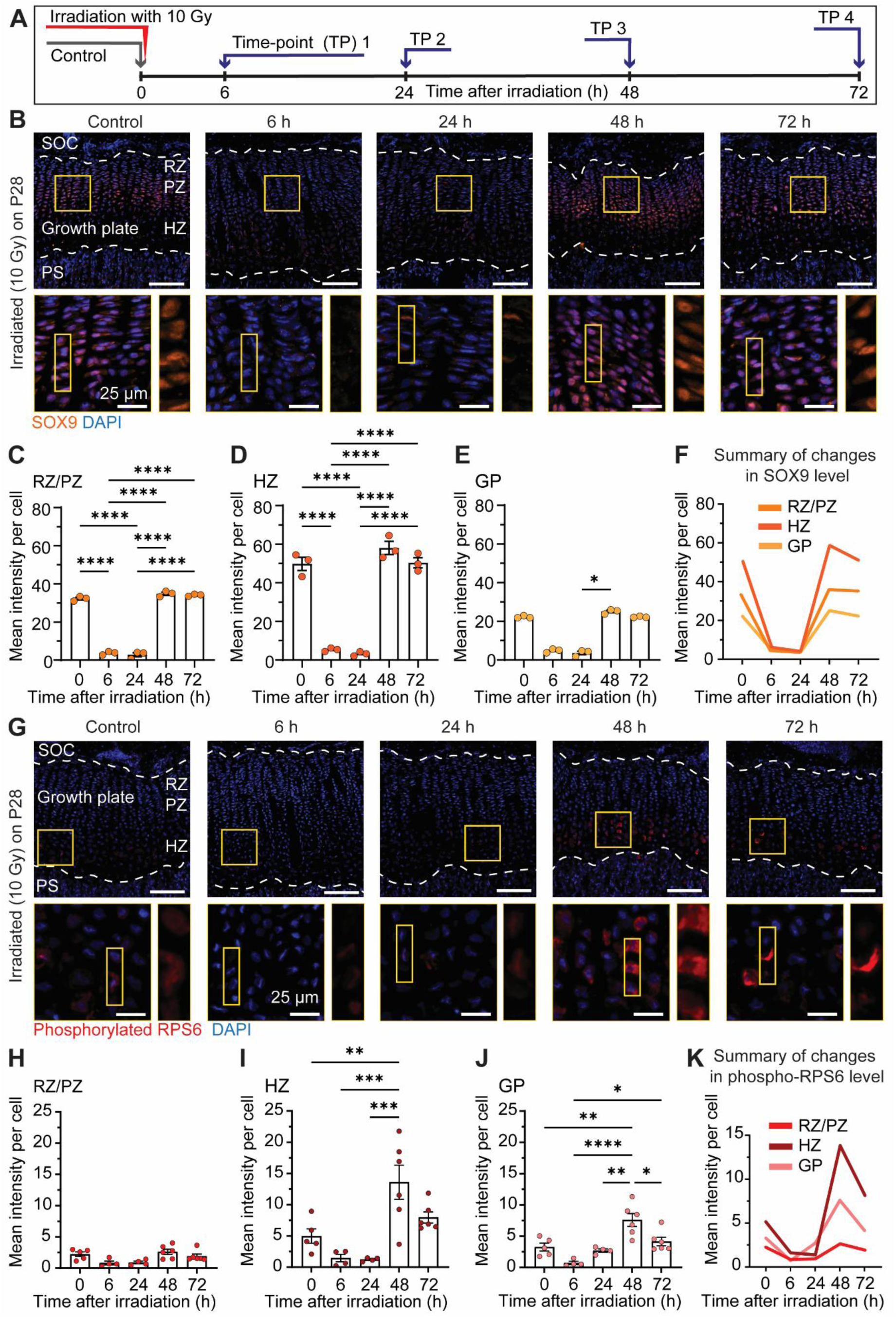
Ionizing irradiation *in vivo* disrupts chondrocyte behaviour. (**A**) Experimental set-up. (**B**) Immunofluorescent images of SOX9 immunostaining, with quantification (**C-E**). **(F)** Merged line graphs to summarize temporal SOX9 alterations. (**G**) Immunofluorescent images of phosphorylated RPS6 in the growth plate at time intervals after 10 Gy irradiation. (**H-J**) Differential quantification of phospho-RPS6 mean intensity per cell in resting and proliferation zones (RZ and PZ), hypertrophic zone (HZ) and entire growth plate (GP) (n≥4 mice). (**K**) Merged line graphs to summarize temporal phospho-RPS6 alterations throughout the growth plate. Scale bars = 100 μm, unless otherwise specified. The secondary ossification center (SOC), growth plate (GP), primary spongiosa (PS), and the resting (RZ), proliferating (PZ) and hypertrophic (HZ) zones are indicated for orientation, and the growth plate is demarcated by dashed lines.

## Results

### Growth plate irradiation disrupts the continuous production of chondrocytes required for bone elongation

Irradiation of the proximal tibial growth plate offers a way to study growth plate injury and repair as mechanisms of growth are similar between growth plates, systemic damage to other organs is mimimized, and late complications can be monitored simply by measuring leg-length discrepancies. Skeletal late complications were induced by applying a single dose of 15 Gy x-ray irradiation to the left proximal tibial growth plate at postnatal day 28 (P28) (Fig. 1B), leading to significant growth retardation 14 days later (Fig. 1C). Growth plate histology was analyzed to explore the acute damage caused by irradiation; within 96 h of irradiation, growth plates were visibly disrupted, with fluctuating height and disorganized hypertrophic zones (Fig. 1D).

Since continuous cell division in the proliferating zone is required to provide hypertrophic chondrocytes and thereby maintain continuous growth [17] (Fig. 1A), we monitored cell dynamics after irradiation injury; after applying ionizing radiation at P28, clonal genetic tracing to label collagen type II alpha 1 chain (Col2a1)-positive chondrocytes was conducted the following day using the R26R-Confetti mouse strain and tissues collected three days later (Fig. 1E). Clones that formed from chondrocytes irradiated 96 hours (h) earlier were positioned throughout the growth plate (Fig. 1F). The clonal sizes were significantly smaller in the irradiated (L) than control (R) growth plates (Fig. 1G), which resulted in significantly more single-celled clones in the irradiated side than contralateral control, while there was a reduction in the number of doublets, triplets and quadruplets in irradiated growth plates (Fig. 1H). The significantly reduced clonal sizes in irradiated growth plates indicated that irradiation disrupts bone elongation by rapidly impairing the production of new cells. These data confirm the efficacy of this approach to irradiate the proximal tibial growth plate.

### Focal tibial irradiation *in vivo* closely models the development of skeletal late complications

To improve the clinical relevance of our irradiation model [20,21], we reduced the irradiation dose to 10 Gy irradiation, corresponding to a biologically effective dose (BED) [18] of 32.37 Gy using an alpha/beta ratio of 4.47 for growing cartilage [19,22], and tested the effect this radiation dose had on tibial growth. Firstly, a focal dose of 10 Gy induced tibial length discrepancy two weeks after irradiation (Fig. 2A). To determine if growth plates were able to form new bone tissue one week after irradiation, we injected a fluorescent dye (calcein) to label newly formed mineral, enabling the quantification of growth rates *in vivo* [23] (Fig. 2B). Newly formed mineralized tissue was deposited across the entire width of irradiated growth plates analyzed (Fig. 2C). However, irradiated growth plates had significantly less (78.65 µm ±13.24) newly formed bone than contralateral control (204.00 µm ±18.64) during the final 48 h before tissue collection (Fig. 2D). Importantly, these results demonstrate that growth plates irradiated with 10 Gy produce new bone tissue within one week of irradiation injury but at reduced rates, which leads to a significant reduction in length several weeks later, closely mimicking the clinical progression of skeletal late complications [13].

### Ionizing radiation causes acute DNA damage throughout the growth plate

Next, we aimed to determine which growth plate cell populations were most susceptible to irradiation damage. To this end, proliferating cells were labelled with the thymidine analogue 5-ethynyl-2’- deoxy uridine (EdU) 4 h prior to irradiation, and tissues collected thirty minutes later (Fig. 2E). A dose of 10 Gy was sufficient to cause DNA double-strand breaks, as assessed by immunofluorescence to γH2A.X variant histone (γH2AX) [24], in chondrocytes within all zones of the growth plate (Fig. 2F). Interestingly, damage was not uniform across the chondrocyte populations, whereby hypertrophic zone chondrocytes had the most DNA lesions (Fig. 2G). Furthermore, irradiated EdU-positive cells did not have significantly more DNA damage than other irradiated growth plate chondrocytes (Fig. 2G). To determine if this DNA damage resulted in apoptosis, we immuno-stained for the cleaved form of caspase 3 (CASP3); less than 1% of chondrocytes were positive for cleaved CASP3 across all time-points (Fig. 2H, I)). These data indicate that while DNA damage occurred throughout the growth plate in response to 10 Gy irradiation, few chondrocytes appeared to undergo cell death.

### Ionizing radiation triggers an acute increase in autophagic flux which returns to normal levels within 48 h

To better understand how growth plates respond to irradiation, the temporal phases of acute injury in this model were assessed over a time-course up to 72 h post-irradiation (Fig. 3A). While SOX9 plays broad roles in various tissues, it has an important role in chondrocytes because it is a transcription factor directly leading to the expression of cartilage matrix proteins [25]. Interestingly, immediately after irradiation, SOX9 expression dropped dramatically in resting/proliferating chondrocytes (Fig. 3B-C), the hypertrophic chondrocytes (Fig. 3B,D) and, indeed, the growth plate as a whole (Fig. 3E). The protein levels of SOX9 were significantly downregulated by 6 h and then returned to detectable levels 48 h after irradiation (Fig. 3F), suggesting fluctuations in chondrocyte functionality. Chondrocyte behavior was also assessed using ribosomal protein S6 (RPS6) as a readout, which is typically phosphorylated in pre-hypertrophic chondrocytes, indicating cells an active transition from proliferative to hypertrophic zones [26]. While there were no significant changes noted in phospho-RPS6 levels in the resting and proliferating chondrocytes specifically (Fig. 3G,H), phospho-RPS6 was markedly reduced in the hypertrophic zone (Fig. 3I), which mirrored the growth plate as a whole (Fig. 3J and K). Together, these results demonstrate that irradiation caused an almost immediate loss of typical behavior to chondrocytes throughout the growth plate but that these features had returned to levels similar to unirradiated controls within 72 h.

Phosphorylated RPS6 is a readout of mammalian target of rapamycin complex 1 (mTORC1) activity [27], a signaling hub that integrates a wide variety of intracellular and extracellular signals, directing anabolism during favorable conditions or otherwise stimulating autophagy during catabolic conditions [28]. Under physiological conditions, autophagy is inhibited in pre-hypertrophic chondrocytes as demonstrated by the accumulation of sequestosome 1 (SQSTM1) and microtubule-associated protein 1 light chain 3 alpha (LC3-II) puncta within these cells [7]. On the other hand, autophagy is an ongoing process in resting and proliferating zone chondrocytes during physiological conditions since conditional knockout mouse models have been used to demonstrate that SQSTM1 accumulates in growth plate chondrocytes lacking autophagy related 5 (Atg5) or Atg7 [7,29]. Autophagy inhibition by mTORC1 hyperactivation induces cell death via lysosomal storage disorders in growth plate chondrocytes [30]. Given the dramatic fluctuations in phospho-RPS6 levels in response to irradiation (Fig. 3K), we decided to assess the levels of autophagy in these mice.

To accurately assess autophagic flux, changes in activity should be monitored over time [31]. Using the time-course presented in Fig. 3A, we immuno-detected SQSTM1, which accumulates in autophagosomes and autolysosomes when autophagy is inhibited [32]. Autophagy is normally inhibited in pre-hypertrophic chondrocytes (Fig. 4A), as indicated by SQSTM1 accumulation. However, SQSTM1 levels decreased 6 and 24 h after irradiation damage, particularly in the hypertrophic zone (Fig. 4A-E), and then accumulated again 48 and 72 h after irradiation. Since lysosome-associated membrane protein 1 (LAMP1) is predominantly associated with lysosomes [33], its levels can indicate changes in the rate of autophagy. Interestingly, opposing effects were seen in different growth plate zones (Fig. 4F), with moderate reduction in the RZ/PZ (where autophagy was already occurring prior to irradiation, Fig. 4G) but a large increase in LAMP1 signal in hypertrophic zone (Fig. 4H-J). Together with SQSTM1 and phospho-RPS6 levels, these results suggest that irradiation stimulates autophagy in the murine growth plate via mTORC1.

**Figure 4.**
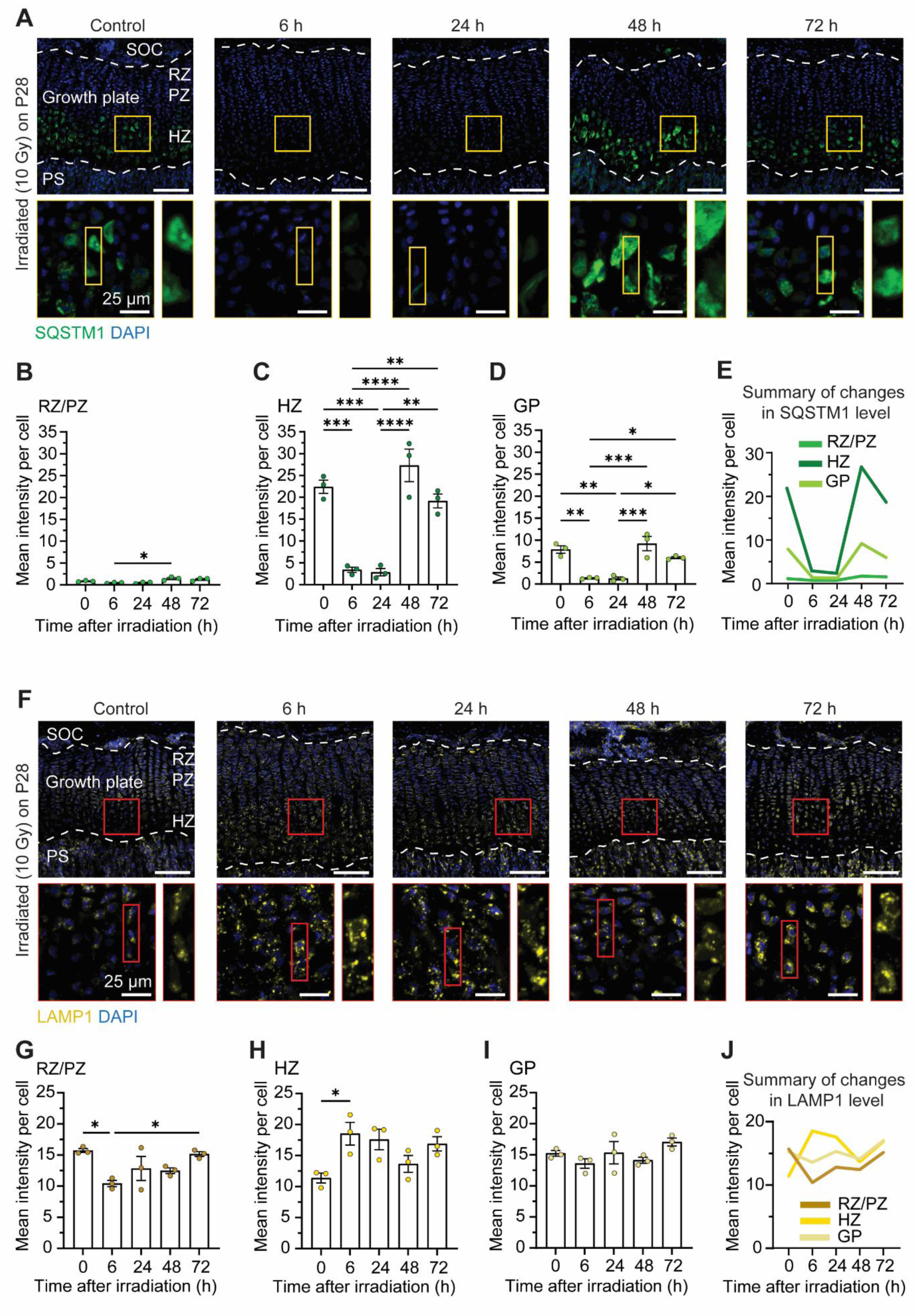
Ionizing irradiation *in vivo* caused an acute elevation in autophagic flux throughout the growth plate. (**A**) Immunofluorescent images of SQSTM1 staining in time intervals after 10 Gy irradiation. (**B-D**) Quantification of temporal SQSTM1 mean intensity per cell in resting and proliferation zones (RZ and PZ), hypertrophic zone (HZ) and entire growth plate (GP; n=3 mice). (**E**) Merged line graphs to summarize temporal SQSTM1 alterations. (**F**) Immunofluorescent images of LAMP1 at time intervals after 10 Gy irradiation. (**G-I**) Quantification of temporal LAMP1 mean intensity per cell in resting and proliferation zones (RZ and PZ), hypertrophic zone (HZ) and entire growth plate (GP) (n=3 mice). (**J**) Merged line graphs to summarize temporal LAMP1 alterations throughout the growth plate. Bars represent mean ± S.E.M., p-value ≤ 0.05. Scale bars = 100 μm, unless otherwise specified. The secondary ossification center (SOC), growth plate (GP), primary spongiosa (PS), and the resting (RZ), proliferating (PZ) and hypertrophic (HZ) zones are indicated for orientation, and the growth plate is demarcated by dashed lines.

**Figure 5.**
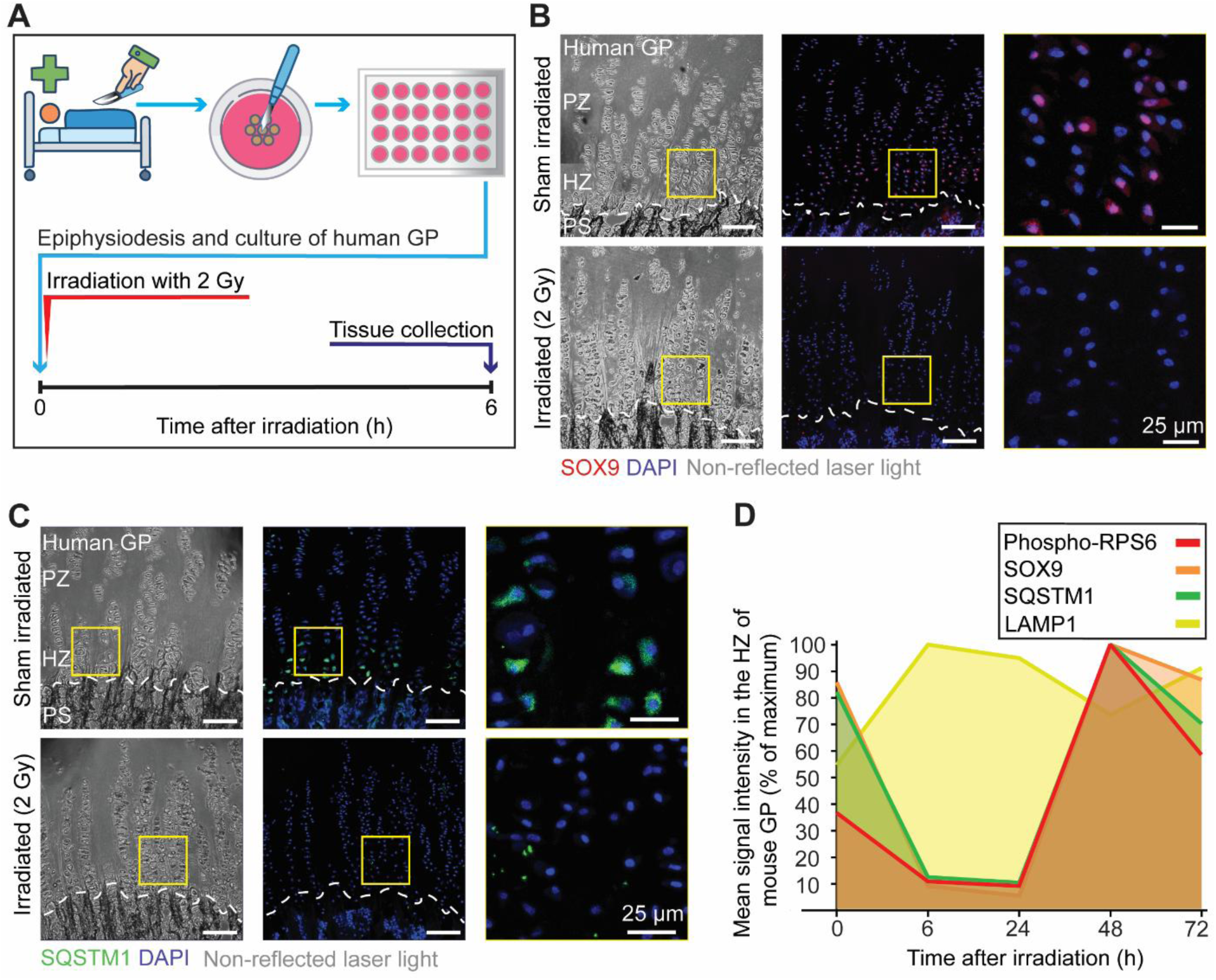
Ionizing irradiation caused an acute loss of SOX9 and SQSTM1 in human growth plate chondrocytes. (**A**) *Ex vivo* experimental set-up: human growth plate slices were irradiated approximately 2-4 h after epiphysiodesis surgery and then cultured for 6 h prior to fixation. Immunofluorescent staining indicated that ionizing irradiation caused a loss of (**B**) SOX9 (representative of n = 3 patients) and (**C**) SQSTM1 (representative of n = 2 patients) throughout the growth plate in comparison with sham-irradiated control slices taken from the same biopsy. (**D**) A summary of the molecular dynamics corresponding to the chondrocyte over the time points after injury with relative intensity of each marker. Scale bars = 100 μm, unless otherwise specified. The growth plate (GP), primary spongiosa (PS), and the proliferating (PZ) and hypertrophic (HZ) zones are indicated for orientation, and the growth plate is demarcated by dashed lines.

**Figure 6.**
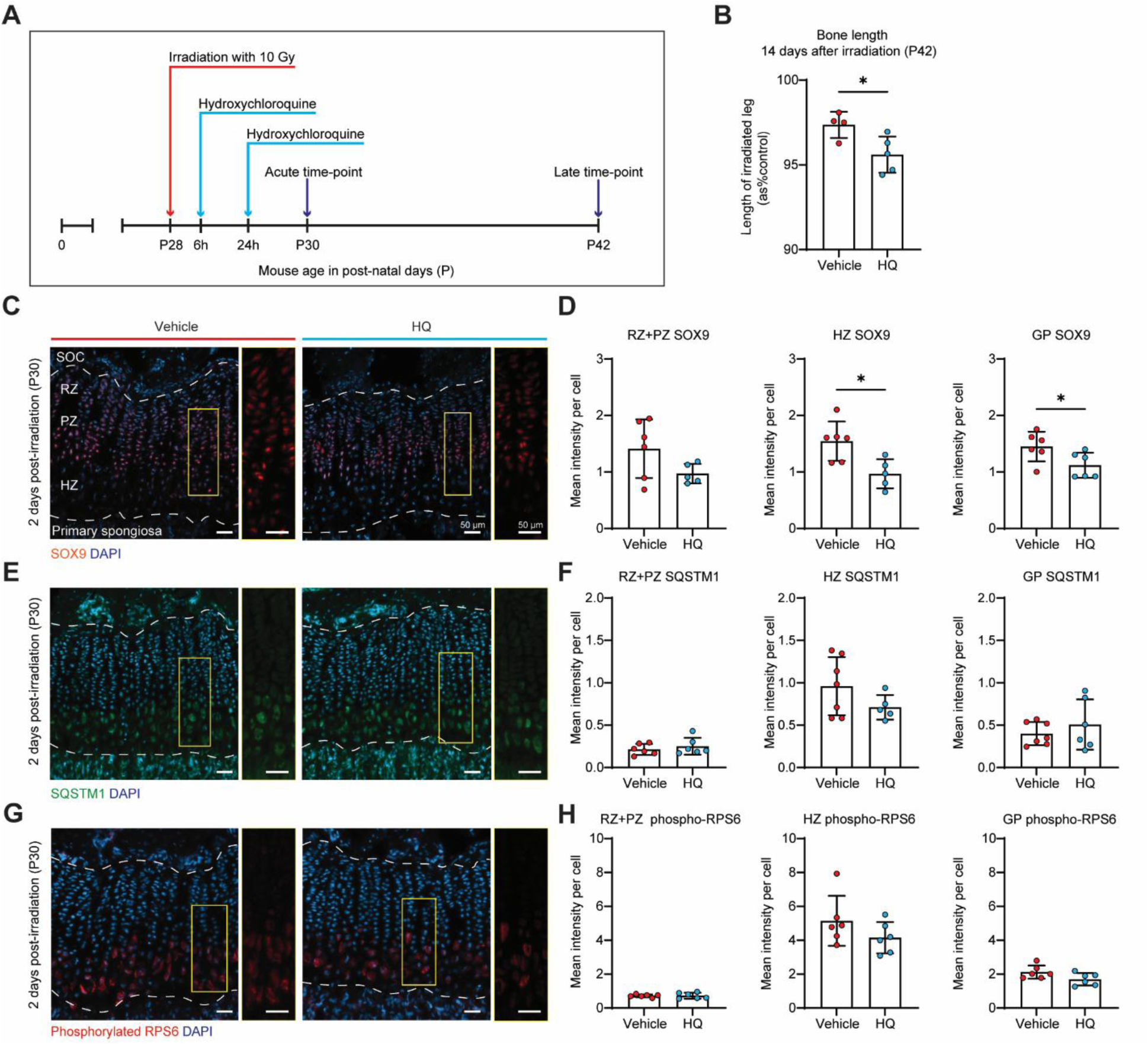
Hydroxychloroquine treatment impairs bone growth recovery and reduces SOX9 expression following irradiation. (**A**) Experimental set-up used to evaluate the effect of autophagy inhibition following irradiation. Analysis was performed at acute (P30, 2 days post-irradiation) or late (P42, 14 days post-irradiation). (**B**) Bone length of the irradiated tibia (L) measured 14 days after irradiation (P42) and expressed as a percentage of the length of the contralateral tibia. (**C**) Confocal microscopy images of immunofluorescence staining of SOX9 in growth plates at P30 following hydroxychloroquine or vehicle treatment, and (**D**) its quantification in the resting and proliferating zones (RZ + PZ), hypertrophic zone (HZ), and the entire growth plate (GP). (**E**) Immunofluorescence staining of SQSTM1 staining at P30, and (**F**) its quantification in RZ + PZ, HZ, and GP. (**G**) Immunofluorescence staining of phospho-RPS6 at P30, and (**H**) its quantification in RZ + PZ, HZ, and GP. The data points on each graph represent individual mice; bars indicate mean ± SEM Abbreviations: secondary ossification centre (SOC), growth plate (GP), primary spongiosa (PS), left (L), right (R), resting zone (RZ), proliferative zone (PZ), hypertrophic zone (HZ).

### Human growth plate chondrocytes respond to ionizing radiation with acute elevation in autophagy and a loss of SOX9

Given the growing interest in autophagy modulation in pediatric oncology (Table 2), we extended our investigation to human tissues. To do this, we obtained human growth plate biopsies and devised a strategy to irradiate them in order to explore the acute effects of irradiation on the chondrocytes (Fig. 5A). As soon as they were prepared for *ex vivo* culture, we administered a single dose of 2 Gy irradiation to mimic the fraction size given clinically [20], and shielded a control section from the same biopsy. Following 6 h in culture, biopsies were fixed and subsequently immuno-stained for SOX9 and SQSTM1. Our results indicated that SOX9 levels plummeted in chondrocytes throughout the human growth plate (Fig. 5B), coinciding with degradation of SQSTM1 in hypertrophic zone (Fig. 5C). These observations strongly indicate that our findings made using our *in vivo* mouse model (Fig. 5D) apply to human chondrocytes embedded within their growth plate cartilage.

### Autophagic flux is required for growth plate regeneration in response to irradiation

Despite the diverse role(s) autophagy plays in different types of cancerous and healthy tissues [9], the use of autophagy modulators is an emerging topic in cancer treatment, including pediatric oncology patients. Autophagy inhibition using, for example, HQ, and activation via mTOR inhibitors (eg. everolimus, sirolimus) are the listed interventions of at least 14 active clinical trials involving children, with more than half in the recruiting/not yet recruiting stage. Specifically, autophagy inhibitors are being tested in breast cancer and specific glioma patients, whereas autophagy activators are being used to treat pediatric patients with solid tumors, metastatic sarcoma, and specific glioma patients, among others (as presented in Table 2). Autophagy modulation during radiotherapy is also being tested in ongoing clinical trials. While most of these interventions focus on the use of autophagy inhibitors to overcome radiation resistance by blocking autophagic processes, others use autophagy inducers to sensitize cancer cells to radiotherapy-induced cell death [34,35].

To assess the functional importance of acute autophagy activation following growth plate irradiation, we inhibited autophagic flux *in vivo* using hydroxychloroquine (HQ) administered at 6 and 24 hours post-irradiation (Fig. 6A), based on the timing of early autophagic changes observed in Fig. 3–4. HQ treatment effectively blocked autophagy, based on SQSTM1 levels, in the irradiated growth plates several hours after the final dose (Fig. 7A-C), and resulted in a significantly greater tibial length discrepancy than irradiation alone (Fig. 6B), indicating impaired recovery. Although autophagy levels had returned to baseline by 48 hours in irradiated mice, SOX9 expression in HQ-treated mice remained low (Fig. 6C, Fig. 7A,D,E), suggesting a failure to re-establish normal chondrocyte function. Importantly, the lengths of the non-irradiated tibiae were unaffected by transient HQ treatment (Fig. 7F), supporting that notion that inhibition of autophagy in the irradiated growth plate mediated this effect. Together, these findings indicate that transient inhibition of autophagy immediately after irradiation exacerbates longitudinal growth defects, supporting a protective role for acute autophagic activation in skeletal recovery.

**Figure 7.**
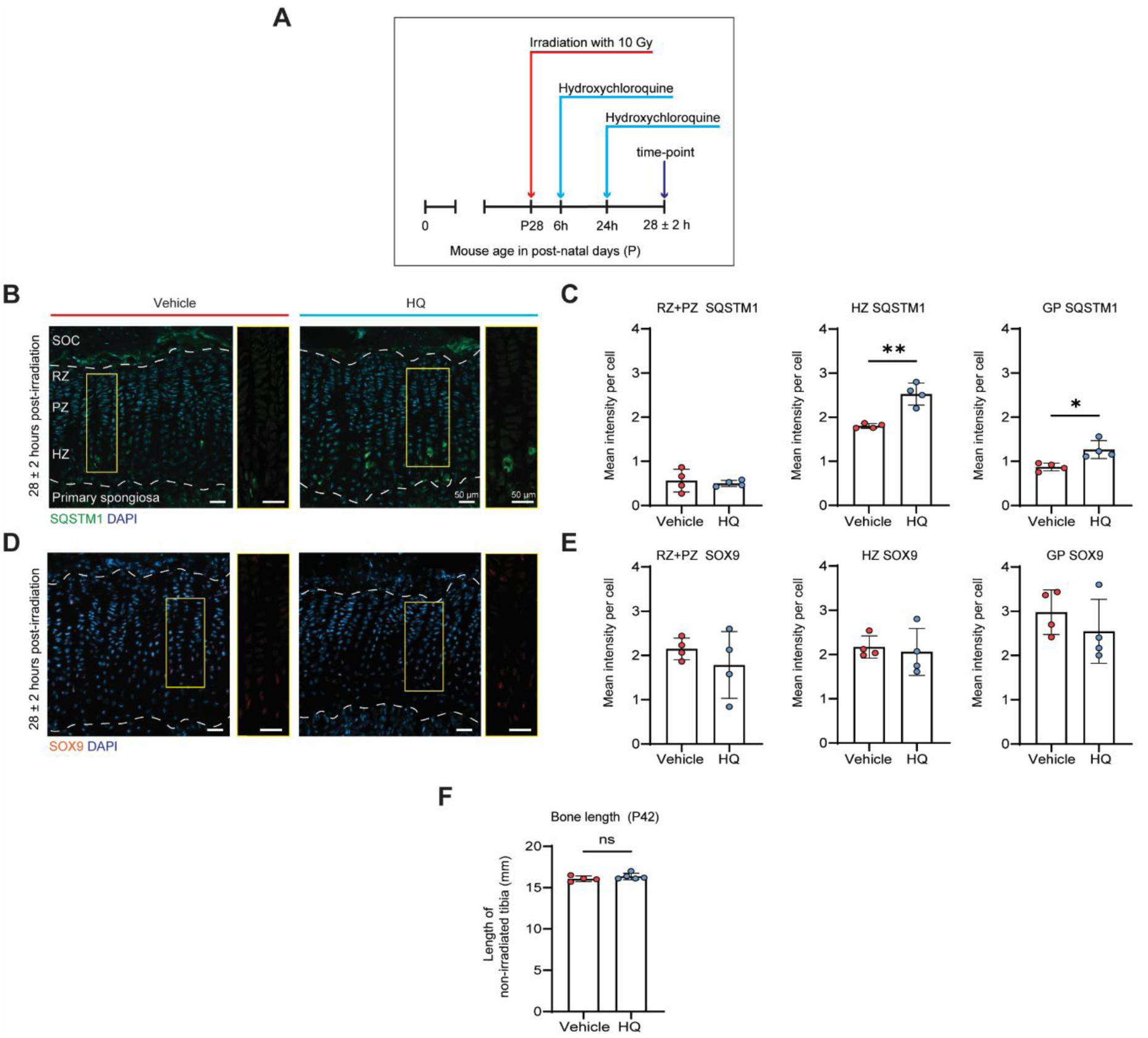
Hydroxychloroquine treatment blocks autophagic flux in the irradiated growth plate. **(A)** Experimental set-up used to evaluate whether the HQ treatment regimen inhibited autophagy within the growth plate following irradiation, as analyzed in B-E. **(B)** Representative immunofluorescence images of SQSTM1 staining. **(C)** Quantification of SQSTM1 mean intensity per cell in resting and proliferation zones (RZ and PZ), hypertrophic zone (HZ) and entire growth plate (GP; n=4 mice). **(D)** Representative immunofluorescence images of SOX9 staining. **(E)** Quantification of SOX9 mean intensity per cell in resting and proliferation zones (RZ and PZ), hypertrophic zone (HZ) and entire growth plate (GP; n=4 mice). **(F)** Length of the non-irradiated tibia, used as a contralateral control (R) in the experiment presented in Fig. 6A, measured 14 days after irradiation (P42). The data points on each graph represent individual mice; bars indicate mean ± SD. Abbreviations: secondary ossification centre (SOC).

By revealing the importance of rapid changes in autophagic flux in response to irradiation, our results suggest that late complications can be affected by modulating autophagy. Hence, monitoring skeletal late complications could be an important outcome measure in future clinical trials of pediatric oncology patients treated with autophagy modulators. Moreover, targeting autophagic flux could be an effective way to treat patients with the aim of preventing skeletal late complications.

## Discussion

Radiotherapy is and will remain is a major part of cancer treatment for children for the foreseeable future whereas there are no preventative interventions against skeletal late complications for patients receiving irradiation to the growth plates, and these patients have limited treatment options once damage has occurred [15]; understanding the biology underpinning radiation-induced late complications is, therefore, necessary to improve quality of survival for generations of pediatric oncology patients [19]. In the present study, we characterized the acute responses of murine and human growth plates to focal irradiation and identified a rapid, transient activation of autophagy as a necessary component of early growth plate recovery. These findings reveal previously unrecognized mechanisms underlying radiation induced skeletal toxicities and highlight autophagy modulation as a potentially modifiable determinant of post-treatment bone growth outcomes.

Despite its importance, research into the molecular mechanisms by which growth plates respond to radiation injury remains limited. The most relevant studies were published from a single group of researchers in the early 2000s: they used microarray analysis of five-week old rat growth plates following 17.5 Gy x-ray irradiation (either as a single dose or in five fractions) and performed laser capture micro-dissection to analyze different growth plate zones between 7 to 16 days post-irradiation [36–38]; while these studies revealed many temporally differentially expressed genes in different zones, a consistent outcome was the aberrant expression of extracellular matrix (ECM) proteins and ECM-remodeling enzymes, and elevated cytokines and growth factors in proliferating and hypertrophic cells; in particular, IGF2 expression in the PZ and CTGF in both the PZ and HZ a week after irradiation [36]. Our findings reveal even earlier changes to chondrocyte behaviour induced by radiation damage that may subsequently influence these transcriptional alterations.

Mechanistically, our study demonstrates that ionizing radiation perturbs mTORC1 activity and induces a rapid, transient increase in autophagic flux in chondrocytes. The acute decline in phosphorylated RPS6 and SQSTM1 levels at 6–24 h, followed by recovery within 48–72 h, indicates dynamic and tightly regulated modulation of autophagy. Similar short lived autophagic responses have been described in irradiated epithelial and neural tissues, where autophagy acts as a cytoprotective mechanism that clears damaged proteins and organelles [39,40]. Importantly, our *ex vivo* human growth plate experiments confirmed that chondrocytes from human tissue mount the same conserved acute response after a single, clinically-relevant, 2 Gy fraction. Our functional experiments with HQ also demonstrate that autophagy activation is not merely a bystander response but is actively involved in continued bone growth after irradiation. Together, these findings support a model in which autophagy activation is a conserved, early adaptive mechanism that helps maintain chondrocyte viability and functional integrity after radiation-induced damage.

To study growth-associated skeletal late complications, we utilised *in vivo* mouse approaches as they provide a reliable way to monitor how these conditions develop over time, as occurs in patients. Mice provide suitable models of human growth plates as the general mechanisms of skeletal growth are highly evolutionarily conserved across extant mammals [41]. We selected the proximal tibial growth plate, as its biology is well described [1,17,25], and its focal irradiation could minimize systemic effects to other organs; thus tibial irradiation offers a simple way to study direct growth plate injury with a clear readout: leg-length discrepancy. Some future refinements could further improve the model; for example, we employed single-dose irradiation delivered in a two-dimensional (2D) configuration, whereas clinical radiotherapy typically involves fractionated dosing delivered via three-dimensional (3D) arc techniques to minimize exposure to surrounding tissues. However, we applied BEDs approximating those used in standard clinical settings; for example, assuming an alpha/beta ratio of 4.47 [19] gave a BED [18] of 32.37 in our set-up (one dose of 10 Gy; used in mouse experiments from Fig. 2 onwards), compared with, for example, a BED of 32.8 applied clinically (1.8 Gy x 13) to growing vertebrae in medulloblastoma patients [42]. Given the acute effects of radiation detected in these mice also occurred in human tissue (at a lower radiation dose of 2 Gy x 1; BED [18] of 2.89 Gy), we consider our results to be relevant to the clinical situation, while there is a broader need for more refined models in this research area.

Clinical trials are underway using both autophagy inhibitors - to overcome radiation resistance by blocking autophagic processes- and autophagy-inducers - to sensitize cancer cells to radiotherapy-induced cell death [34,35]. While most interventions focus on the use of autophagy inhibitors, the majority of clinical studies have also focused on cancers in adults [34]. The interest in this area has led to the modulating autophagy in childhood cancers in pre-clinical studies [43–45]. Additional studies conclude that autophagy modulators could be reasonable targets in pediatric patients [43,44] and has resulted in several ongoing clinical trials in pediatric oncology patients [45]. In this context, our results are highly relevant and extremely timely.

Given that growth plates are commonly exposed to irradiation in addition to the targeted organs, the potential for skeletal late complications should be a consideration when developing autophagy-modulating strategies. Our findings not only reveal key molecular mechanisms underlying acute growth plate regeneration following radiation injury, but also highlight potential therapeutic opportunities to prevent or mitigate radiation-induced skeletal late complications. The rapid alterations in autophagic flux observed in our experiments underscore the importance of timing, as interventions administered at different phases may produce markedly different outcomes in irradiated growth plates. This complexity is further compounded by the fact that distinct zones of the growth plate exhibit different basal levels of autophagy [7,29], which could influence their responsiveness to treatment. Therefore, whether therapeutic strategies are explicitly designed to modulate autophagy within the growth plate or applied more generally, these observations emphasize the need for careful consideration of timing, dosing, and tissue-specific effects when using autophagy-modulating agents in children receiving radiotherapy.

## Conclusions

We provide insights into the acute response of growth plate chondrocytes to ionizing irradiation injury, characterized by elevated autophagic flux, which allows them to maintain limited growth and partially recover from radiation-induced damage, but ultimately leads to irradiation-induced skeletal late complications. These findings underscore the necessity for refined therapeutic strategies to mitigate the adverse effects of radiotherapy on skeletal growth in children, paving the way for future research focused on enhancing regenerative outcomes through targeted autophagy interventions. Ultimately, understanding these processes not only contributes to the field of regenerative medicine but also holds promise for improving the long-term health and quality of life for pediatric cancer survivors facing the consequences of ionizing radiation exposure.

## Acknowledgements

Irradiation experiments were performed at the X-ray Irradiation Core Facility (Karolinska Institutet). We would like to acknowledge Dr. Sofia Skyttner for substantial technical and theoretical support relating to radiation dosimetry and experimental setup. We also wish to thank Dr. Adamantia Fragkopoulou and Dr. Ahmed Osman for additional technical support, and Prof. Lev Novikov for his generous support of YMA. PTN was financially supported by the Barncancerfonden (PR2019-0049, PR2022-0035), Cancerfonden (23 2739 Pj), and the European Science Foundation. This project was partially supported by the Fight Kids Cancer Funding Programme, supported by Imagine for Margo, Foundation KickCancer, Foundatioun Kriibskrank Kanner, CRIS Cancer Foundation, and Children Cancer-free Foundation.

